# Nanopore long-read transcriptome data of fungal pathogen of chalkbrood disease, *Ascosphaera apis*

**DOI:** 10.1101/2020.03.12.989863

**Authors:** Yu Du, Huazhi Chen, Jie Wang, Zhiwei Zhu, Cuiling Xiong, Yanzhen Zheng, Dafu Chen, Rui Guo

**Affiliations:** College of Animal Sciences (College of Bee Science), Fujian Agriculture and Forestry University, Fuzhou 350002, China

**Keywords:** Nanopore, long-read transcriptome, chalkbrood, *Ascosphaera apis*, honeybee

## Abstract

*Ascosphaera apis* is a fungal pathogen that exclusively infects honeybee larvae, leading to chalkbrood disease, which damages the number of adult honeybees and colony productivity. In this article, *A. apis* mecylia and spores were respectively purified followed by Oxford Nanopore sequencing via PromethION platform. In total, 6,321,704 and 6,259,727 raw reads were generated from Aam and Aas, with a length distribution among 1 kb~10 kb. The quality (Q) scores of majority of raw reads were Q9 (Aam) and Q11 (Aas). Additionally, 5,669,436 and 6,233,159 clean reads were gained, among them 79.32% and 79.62% were identified as being full-length. The lengths of redundant reads-removed full-length transcripts were among 1 kb~8 kb and 1 kb~9 kb, and most abundant length for both was 1 kb. Furthermore, the length of redundant transcripts-removed clean reads was ranged from 1 kb~7 kb, with the largest group of 1 kb. The data reported here provides a beneficial genetic resource for improving genome and transcriptome annotations of *A. apis* and for exploring alternative splicing and polyadenylation of *A. apis* mRNAs.

**Value of the result:**

- Current dataset enables better understanding of the complexity of *A. apis* transcriptome.
- The long-read transcriptome data can be used to identify of genes and transcripts associated with *A. apis* infection mechanism.
- The accessible data provides full-length transcripts for improving gene structure and functional annotation of *A. apis* transcriptome.
- This dataset could be utilized for investigation of alternative splicing and polyadenylation of *A. apis* mRNAs.

## 1. Data description

The long-read transcriptome data presented here were from purified mycelium (Aam) and purified spores (Aas) of *A. apis* pure culture under laboratory condition. Totally, 6,321,704 and 6,259,727 raw reads were generated from Aam and Aas, with an average read length of 992 bp and 1047 bp and an N50 of 1094 bp and 1157 bp, respectively (**Table 1**). For both Aam and Aas, the length of raw reads was distributed among 1 kb~10 kb, and the most abundant length was 1 kb (**Figure 1 A,B**). In addition, the quality (Q) scores of majority of raw reads were Q9 (Aam) and Q11 (Aas) (**Figure 2 A,B**). After discarding rRNA, 5,669,436 and 6,233,159 clean reads were gained, and among them 79.32% and 79.62% were identified as being full-length (**Table 2**). The length of full-length clean reads from Aam was distributed among 1 kb~8 kb, while that from Aas was distributed among 1 kb~9 kb (**Figure 3 A,B**); the most abundant lengths for both were 1 kb (**Figure 3 A,B**). After removing redundant reads, the lengths of remaining full-length transcripts of both Aam and Aas were ranged from 1 kb to 7 kb, with the largest group of 1 kb (**Figure 4 A,B**).

**Table 1.** Overview of Nanopore raw reads

| Sample | Reads number | Base number | N50<br>(bp) | Mean<br>length<br>(bp) | Maximum length<br>(bp) |
| --- | --- | --- | --- | --- | --- |
| Aam | 6,321,704 | 6,271,320,854 | 1094 | 992 | 9421 |
| Aas | 6,259,727 | 6,553,996,867 | 1157 | 1047 | 13,060 |

**Table 2.** Summary of full-length clean reads

| Sample | Number of<br>clean reads | Number of full-length<br>clean reads | Percentage of full-length<br>clean reads |
| --- | --- | --- | --- |
| Aam | 5,669,436 | 4,497,102 | 79.32% |
| Aas | 6,233,159 | 4,963,101 | 79.62% |

**Figure 1.**
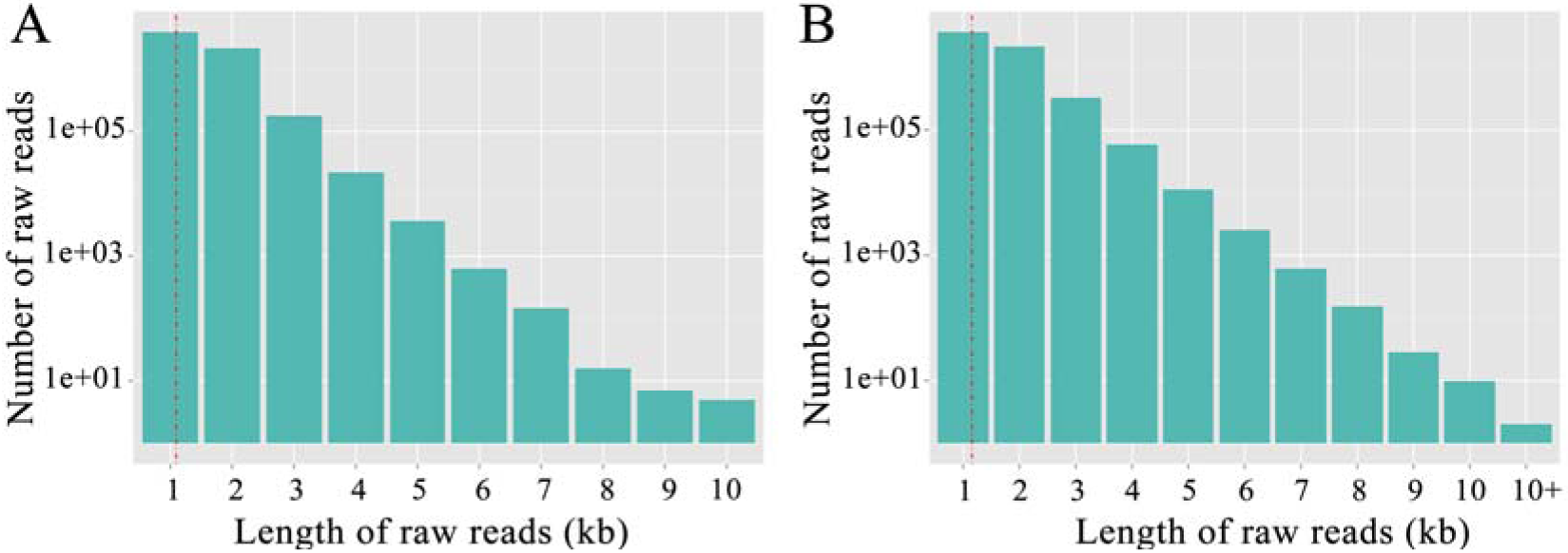
Length distribution of raw reads generated from Nanopore sequencing. (A) Raw reads of Aam; (B) Raw reads of Aas.

**Figure 2.**
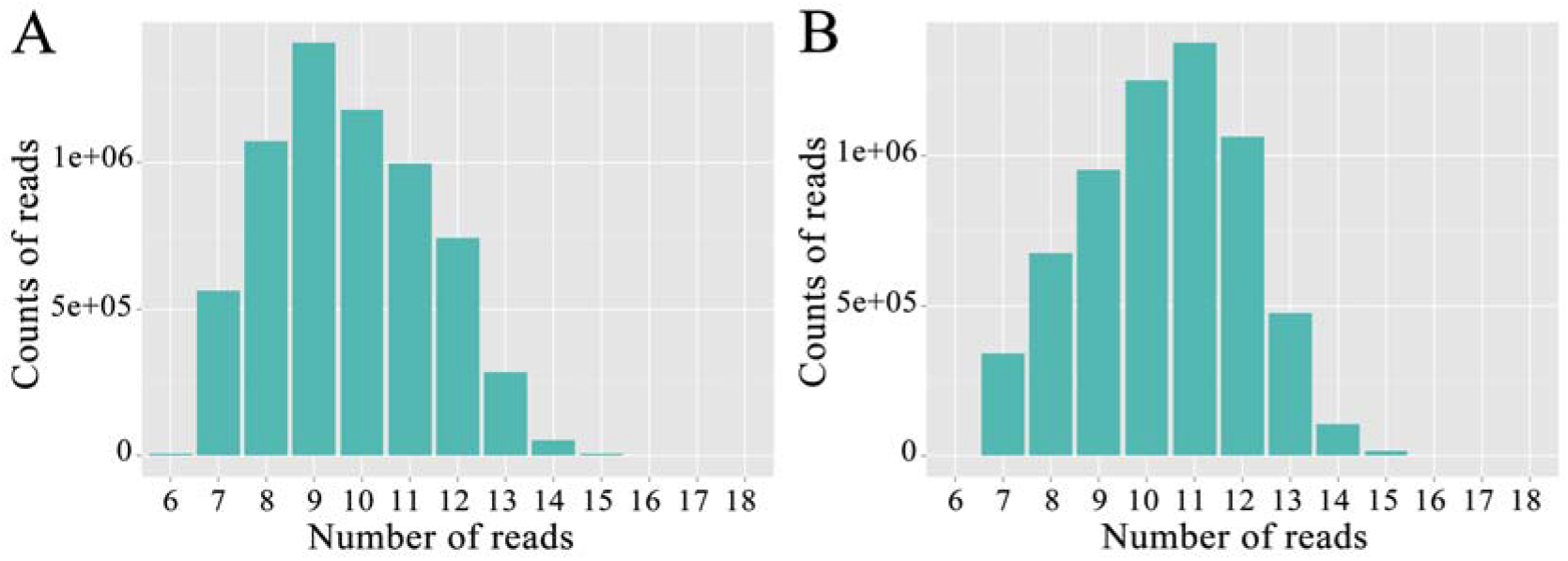
Quality distribution of raw reads derived from Nanopore sequencing. (A) Raw reads of Aam; (B) Raw reads of Aas.

**Figure 3.**
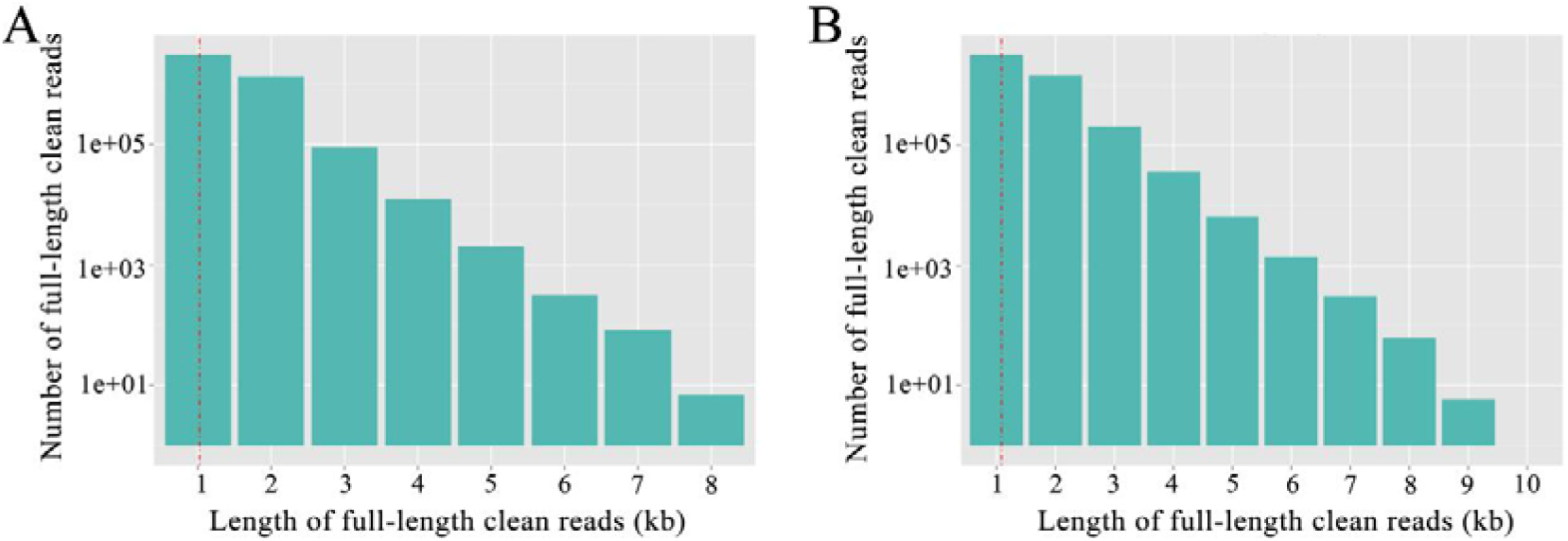
Length distribution of full-length clean reads. (A) Full-length clean reads of Aam; (B) Full-length clean reads of Aas.

**Figure 4.**
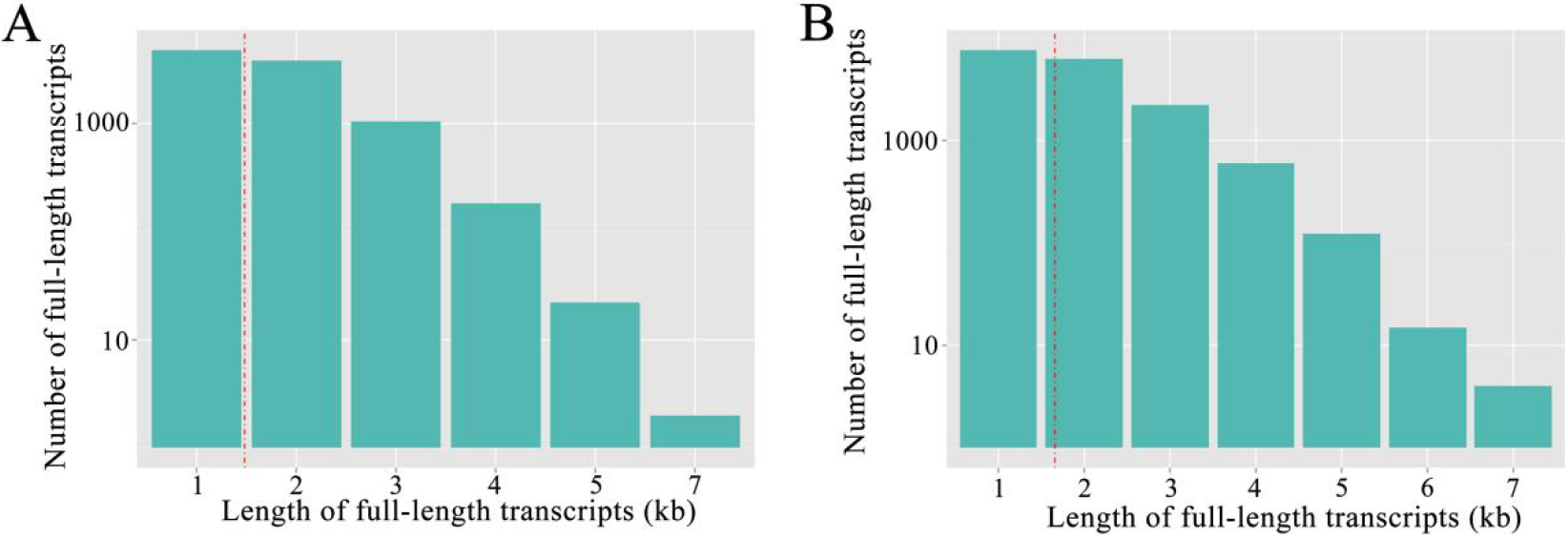
Length distribution of full-length transcripts after removing redundant reads. (A) Redundant reads-removed full-length transcripts of Aam; (B) Redundant reads-removed full-length transcripts of Aas.

## 2. Experimental Design, Materials, and Methods

### 2.1 Preparation of A. apis mycelium and spore sample

*A. apis* was previously isolated from a fresh chalkbrood mummy of *Apis mellifera ligustica* larva[1] and kept at the Honeybee Protection Laboratory of the College of Animal Sciences (College of Bee Science) in Fujian Agriculture and Forestry University. According to the previously described method[2], *A. apis* was cultured at 33±0.5 °C on plates of Potato-Dextrose Agar (PDA) medium. Mycelia and spores were respectively purified following the method previously developed by Jensen et al.[2] with some modifications: (1) mycelia on the top of the medium were collected and purified one week after culturing; (2) a layer of mycelia above black ascomata were seriously removed with a sterile inoculating loop; (3) ascomata were harvested followed by purification of spores via differential centrifugation and microscopic observation; (4) *A. apis* mycelium and spores were immediately frozen in liquid nitrogen and stored at −80 °C until third-generation sequencing.

### 2.2 RNA isolation, cDNA library construction and Nanopore sequencing

Firstly, total RNA of *A. apis* mycelium sample and spore sample were respectively isolated using TRizol reagent (Thermo Fisher, Shanghai, China) on dry ice and then reversely transcribed using Maxima H Minus Reverse Transcriptase, followed by purification and concentration on a AMPure XP beads. Secondly, cDNA libraries were constructed from 50 ng total RNA with a 14 cycles PCR using the cDNA-PCR Sequencing Kit (SQK-PCS109) and PCR Barcoding Kit (SQK-PBK004) following the manufacturers protocol (Oxford Nanopore Technologies Ltd, Oxford, UK). Thirdly, the aforementioned cDNA libraries were sequenced by Biomarker Technologies (Beijing, China) utilizing PromethION system (Oxford Nanopore Technologies Ltd, Oxford, UK).

### 2.3 Long reads processing

(1) Raw reads were filtered with minimum average read Q score=7 and minimum read length=500bp; ribosomal RNA were discarded after mapping to rRNA database. (2) Full-length non-chemiric (FLNC) transcripts were determined by searching for primer at both ends of reads. (3) Clusters of FLNC transcripts were obtained after mapping to reference genome with mimimap2 [3], and consensus isoforms were gained after polishing within each cluster by pinfish (https://github.com/nanoporetech/pinfish); consensus sequences were mapped to the reference genome (assembly AAP 1.0) using minimap2. (4) Mapped reads were further collapsed by cDNA_Cupcake package (https://github.com/Magdoll/cDNA_Cupcake) with min-coverage=85% and min-identity=90%; 5’ difference was not considered when collapsing redundant transcripts.

## Acknowledgments

This research was supported by the Earmarked Fund for China Agriculture Research System (No. CARS-44-KXJ7), the Science and Technology Planning Project of Fujian Province (No. 2018J05042), the Teaching and Scientific Research Fund of Education Department of Fujian Province (No. JAT170158), the Outstanding Scientific Research Manpower Fund of Fujian Agriculture and Forestry University (No. xjq201814), and the Scientific and Technical Innovation Fund of Fujian Agriculture and Forestry University (No. CXZX2017342, No. CXZX2017343).

